# The Potential of *Capsicum annum* Extracts to Prevent *Lactococcosis* in Tilapia

**DOI:** 10.1101/2020.06.09.142406

**Authors:** Kunda Ndashe, Stellah Ngh’ake, Emelda Pola, Emmanuel Masautso Sakala, Emmanuel Kabwali, Ladslav Moonga, Alexander Shula Kefi, Bernard Mudenda Hang’ombe

## Abstract

The capsaicin was extracted in-house from locally purchased chili pepper (*Capsicum annum*) using the conventional solvent extraction method. Varying concentrations of capsaicin were mixed with Lactococcus garvieae each and inoculated on Mueller Hinton agar to determine the minimum bactericidal concentration. Four groups of 100 fish each were injected with either 1) Capsaicin, 2) bacteria and capsaicin, 3) bacteria and 4) normal saline (negative control). The fish were observed for 7 days post treatment and experiment was repeated three times. Protection against infection was measured by the lack of clinical disease and survivability of the fish during period of experimentation. The minimum bactericidal concentration of capsaicin on growth of *Lactococcus garvieae* was 0.1967mg/ml. Significantly, more fish in the bacteria and capsaicin group survived (p<0.0001) compared to those injected with bacteria only. The findings suggest that chili pepper extract can prevent *Lactococcus garvieae* infection in tilapia.

## 1. Introduction

Globally, there has been an increasing demand for farmed fish in the past decade due to stagnation and eventual decrease of catches in capture fisheries. The aquaculture industry plays an important role in meeting fish demand deficits and also contributes to employment, income and the national economy(Tran et al., 2019). The growth of the aquaculture industry has resulted into the increase of disease outbreaks in fish farms. Some of bacteria that have been identified to cause disease outbreaks in tilapia are *Aeromonas hydrophila, Streptococcus agalactiae, Streptococcus iniae*, and *Lactococcus garvieae*(Amal et al., 2018; Bwalya et al., 2020; Harikrishnan and Balasundaram, 2005). The common treatment regime that may be used for bacterial infection are antibiotics. Unfortunately, indiscriminate use of antibiotics leads to resistance as documented in several bacterial diseases in tilapia (Harikrishnan and Balasundaram, 2005).

Aquaculture production depends on the use of antibiotics to control bacterial disease outbreaks although regulations on antimicrobial administration differs between countries. Currently, measures to protect farmed fish from pathogenic bacteria without using antibiotics are under investigation. For example, probiotic and ethnoveterinary medicines are currently being used in livestock production and are proving to be an effective alternative to antibiotic use (Maroyi, 2012). Ethnoveterinary medicines are products of plant or animal origins and they provide low-cost alternatives to allopathic drugs. They have been used successfully in cattle, sheep, goat and poultry production in Africa, south America and Asia (Confessor et al., 2009; Khan and Hanif, 2006; Maroyi, 2012; Shen et al., 2010). A wide range of microorganisms including Gram-negative and Gram-positive bacteria have been used as probiotics in aquaculture such as *Lactobacillus, Lactococcus, Leuconostoc, Enterococcus, Carnobacterium, Shewanella, Bacillus, Aeromonas, Vibrio, Enterobacter, Pseudomonas, Clostridium*, and *Saccharomyces* species (Cruz et al., 2012). Probiotics supplementation in diets of tilapia have shown success in preventing disease outbreaks of bacterial agents (Hai, 2015).

Hot red pepper (*Capsicum annum*)is one of the most important herbs, which is widely used in human feed world-wide and it belongs to *Solanaceae* family, genus *Capsicum* which are the most heavily and frequently consumed as spices throughout the world (Kobata et al., 1998). Hot red pepper has been used in curative and prophylactic treatment of many terrestrial animal diseases(Akhtar et al., 2000; El-Deek et al., 2012; Shen et al., 2010). Capsinoids such as capsaicin present in hot red peppers have shown antimicrobial activities against disease caused by bacteria and plays prominent role as an immunity booster and anti-inflammatory agent (Kobata et al., 1998; Omolo et al., 2014). In study by El-Deek (2012). *Capsicum annum* supplementation had a significant positive effect on body weight gain, when was used as a supplement to oxytetracycline in broiler production (El-Deek et al., 2012).

So far research has shown the effect of Capsinoids on bacteria and it positive effects when used on terrestrial animals but little is known on its effect on bacterial diseases in aquatic animals. The aim of this study was to determine the efficacy of *Capsicum annum* extracts in preventing *Lactococcus garvieae* infection in three-spot tilapia (*Oreochromis andersonii)* as an alternative to antibiotics.

## 2. Materials and Methods

The study was performed in accordance with the guidelines of the National Health Research Ethics Committee of Zambia and the protocol was approved by the Excellence in Research Ethics and Science (ERES) Converge, a private Research Ethics Board (Protocol Number: 2019-Aug-024). Minimum stress was exerted on the fish during handling and sampling. The fish were sampled and drawn from the experiment (euthanized) at the first appearance of clinical signs. The clinical signs of the disease included lethargy, disorientation, aberrant swimming. The live fish was euthanized with an overdose of clove solution at a concentration of 5g/l of water.

### 2.1. Fish

A total of 410 healthy three spot tilapia (*Oreochromis andersonii*) with mean weight of 100 ± 10g were collected from a Fish Farm in Lusaka district. The aquaculture facility had no previous history of disease outbreaks and the subjects were transported by road in oxygenated bags to the University of Zambia, School of Veterinary Medicine wet-lab. The fish were kept in 500 L tanks supplied with flow-through de-chlorinated water and aerated using stone bubblers. They were allowed to acclimatize for 10 days prior to commencement of the experiment. The fish were fed daily on commercial pellets at a rate of 3% body weight. Daily water temperature averaged 20 ±2°C, mean daily dissolved oxygen was 7.9 ± 2 mg/L and pH was 7 ± 0.2.

### 2.2. Preparation of Capsaicin (*Capsicum annum* extract)

*Capsicum annum* (chili pepper) weighing 3kg was purchased from a supermarket in Lusaka for this study. No effort was made to identify specific origins of chili or to differentiate batches after purchase. The chili was transported to the University of Zambia, School of Veterinary Medicine for sorting and removal of seeds using scalpel blade. The remaining pitch and peel of the chili pepper was sun-dried for 5 days to allow loss of water and concentration of capsaicinoids. The dried chili pepper was then blended to fine power with final weight of 0.3kg. Extraction of the capsaicinoids was done according to (Kurian and Starks, 2002). Briefly, The dried pepper sample was placed in the blender with 50mL methanol and subjected to high-speed mixing for 5 min. The slurry (due to the greatly reduced mass of pulp from the dried pepper) was filtered directly through double Whatman GF/A glass fiber filters at a much faster rate than undried pulp. The smaller quantity of dried pulp residue was washed clean of adsorbed capsaicinoids with smaller volumes of added methanol. Collected pulp was rinsed thoroughly and discarded, and the filtrate volume was adjusted to 100 mL with methanol. The extracted sample was transported to Zambia Bureau of Standards (ZABS), for determination of Capsaicin levels in the dried *Capsicum annum* using High Performance Liquid Chromatography according to (Usman et al., 2014).

### 2.3. Growth inhibition test and Minimal Bactericidal Concentration

The *Capsicum annum* extract (capsaicin) was serially diluted (1:10) from the stock solution to 5 dilutions. *Lactococcus garvieae* cultures were grown on Muller-Hinton agar overnight then diluted with sterilized normal saline to obtain 0.5 (Marc-Farland) MCF of (Densimat ref. 99535 A version, Biomerieux). An equal volume of 0.5ml of the *Lactococcus garvieae* suspension and the capsaicin dilution were mixed and immediately inoculated on Muller-Hinton agar using the spread plate method. The inoculated plates were then stored at 25°C overnight. The inoculated plates were examined and bacterial colonies counted every 24hours for 5 day.

The minimum bactericidal concentration (MBC) was determined by culture of Mueller-Hinton agar plates where no visible bacterial growth. The MBC was defined as the lowest concentration (mg/ml) of the extract in agar plate showing no visible bacterial growth. All these tests were repeated three times.

### 2.4. The Challenge Experiment

Prior to the start of the challenge experiment, 10 fish were sacrificed, sampled and tested for the presence or absence of bacterial infections by bacteria isolation. The fish (n = 400) were divided into in 4 groups (Control, bacteria, bacteria and capsaicin and capsaicin only). The control fish were injected with normal saline: bacteria group were injected with *Lactococcus garvieae* only: the bacteria and capsaicin group were treated with *Lactococcus garvieae* and Capsaicin and Capsaicin group were injected with capsaicin only. The total number of fish per group was 100 individuals. Each group was placed in a separate tank (T1-T4). All fish were injected intraperitoneally with 0.1ml of normal saline, *Lactococcus garvieae* suspension (9.6 ×10^5^ cfu bacteria/fish) only, *Lactococcus garvieae* suspension (9.6 ×10^5^ cfu bacteria/fish) and Capsaicin (minimum bactericidal concentration) and capsaicin (minimum bactericidal concentration) only for treatment groups T1 to T4, respectively. The fish in all the treatment groups were observed for clinical signs of disease and mortality for 7days after the initial treatments.

### 2.5. Statistical Analysis

Statistical analysis was done by determining the Fischer’s exact and Chi square tests were used to determine independence or association between the treatment groups and outcomes of treatment measured by survivability of the fish post treatment with the help of Graphpad Prism 8.0 (www.graphpad.com).

## 3. Results

The concentration of capsaicin in the *Capsicum annum* extracts was 19.67mg/g and when diluted to stock solution was 1.967 mg/ml (Table 1).

**Table 1:** Concentration of capsaicin (mg/ml) in the dilutions of the *Capsicum annum* extract

|  | Initial extract (mg/g) | Stock Solution (100ml)-mg/ml | Concentration in liquid (mg/ml) after serial dilutions |  |  |  |
| --- | --- | --- | --- | --- | --- | --- |
|  |  |  | 1:10<sup>1</sup> dilution | 1:10<sup>2</sup> dilution | 1:10<sup>3</sup> dilution | 10<sup>4</sup> dilution |
| **Capsaicin Concentration** | 19.67 | 1.967 | 0.1967 | 0.01967 | 0.001967 | 0.0001967 |

The minimum bactericidal concentration of *Capsicum annum* extract that showed no growth of *Lactococcus garvieae* was 0.1967mg/ml (Table 2). *Lactococcus garvieae* did not show any susceptibility to *Capsicum annum* extract < 0.01967mg/ml (Table 2).

**Table 2:** Colony forming unit of *Lactococcus garvieae* per 0.1ml with different concentrations of capsaicin

| Day of incubation | Treatment 1 Capsaicin (0.001967mg/ml) | Treatment 2 Capsaicin (0.01967mg/ml) | Treatment 3 Capsaicin (0.1967mg/ml) | Treatment 4 Capsaicin (1.967mg/ml) |
| --- | --- | --- | --- | --- |
| Day 1 | 25 | 0 | 0 | 0 |
| Day 2 | 90 | 12 | 0 | 0 |
| Day 3 | 120 | 50 | 0 | 0 |
| Day 4 | 120 | 142 | 0 | 0 |
| Day 5 | 120 | 142 | 0 | 0 |

The results of the fish in the 4 treatment groups showed a higher survival rates of the fish in the *Lactococcus garvieae* and Capsaicin (80%)and control (95%) groups (Figure 1). The *Lactococcus garvieae* only group showed the least survival rate (0%) among all the treatment groups. During the second day post treatment, mortality (20%) was recorded in the *Lactococcus garvieae* and Capsaicin group after which it stabilised to the end of the experiment. In the Capsaicin only group, mortality (20%) was only recorded on the fourth day post treatment. There was, however, a significant difference (p<0.0001) in the survivability of fish between groups.

**Figure 1:**
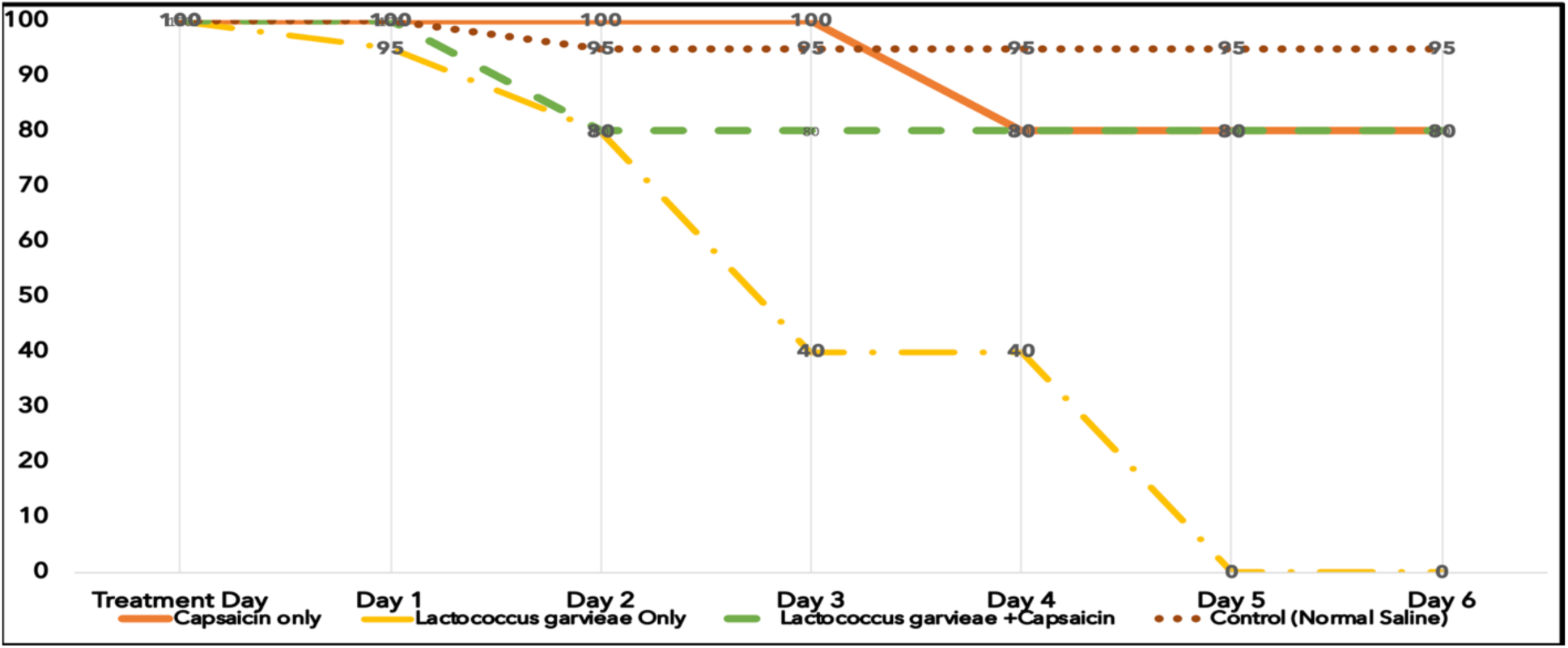
survivability of the fish in the different treatment groups over period of 6 days post treatment

## 4. Discussion

In this study, *Capsicum annum* extract showed antibacterial activities against *Lactococcus garvieae*. This study showed that capsaicin concentration in the local chili pepper was 19.67mg/g and the 0.1967mg/ml was the MBC to inhibit on the growth of *Lactococcus garvieae* on culture media. The MBC of capsaicin in this study is conformity with other studies that indicated the minimum inhibitory concentration (MIC) of capsaicin to range from 0.1 to 0.2mg/ml (Fieira et al., 2013; Marini et al., 2015). Marini and colleagues (2015) demonstrated that the capsaicin minimum inhibitory concentrations (MICs) of erythromycin-resistant and erythromycin-susceptible strains were prevalently in a narrow range (0.064–0.128 mg/mL), where the most common MIC was 0.128 mg/mL and the highest MIC was exhibited at 0.512 mg/ml (Marini et al., 2015). In a study by Careaga and colleagues (2003), they reported a varying degrees of inhibition against *Salmonella typhimurium* inoculated in raw beef meat (Careaga et al., 2003). Other researcher reported that Capsicum extracts had an inhibitory effect on *Salmonella typhimurium, Staphylococcus aureus, Listeria monocytogenes*, and *Bacillus cereus* (Koffi-Nevry et al., 2012).

Prophylactic administration of *Capsicum annum extract* to tilapia infected with *Lactococcus garvieae* showed very good outcome. Significantly, more (80%) fish in the capsaicin and bacteria treatment group survived to the end of the experiment as compared to the *Lactococcus garvieae* only which all the fish died (p<0.0001). The results demonstrated the possibility of *Capsicum annum* extract having an effect on the pathogenesis of *Lactococcus garvieae* infection in the fish. In other trials with *Capsicum annum* on livestock, a study in poultry revealed that it was effective in the control against Newcastle Disease in broilers (Hanif et al., 2012). *Capsicum annum* has been used as an ethnoveterinary medicine for prophylactic treatment of diseases of terrestrial animals mainly domestic livestock (Cichewicz and Thorpe, 1996; Guèye, 2002). With the increasing incidence of antimicrobial resistance to commercially available antibiotics such as oxytetracycline, the use ethnoveterinary medicines like *Capsicum annum* seems a more sustainable method to control livestock disease. And since aquaculture farms have been considered to significantly facilitate exchange of antibiotic resistance genes through contamination of diversity of aquatic organisms with resistant microorganisms (Watts et al., 2017), the use of *Capsicum annum* as an antimicrobial agent may offer solution to curb antimicrobial resistance in the aquaculture sector.

## 5. Conclusion

The results of the present study suggests that three spot tilapia can be treated with *Capsicum annum* extract and protected against *Lactococcus garvieae* by intra peritoneal injection. However, because of the route of injection for challenge and the relatively small number per group of the fish, additional studies are required to confirm with greater certainty the findings of the present study. Further studies should be undertaken to determine the possibility of the administration of *Capsicum annum* extracts through feed and its effects on other aquatic microorganisms.

## 6. Acknowledgements

The work was supported by the financial contributions of the authors. Further gratitude is extended to Professor Bernard Hang’ombe for making available the bacteriology and wet laboratory to conduct the experiments.

## References

Akhtar, M.S., Iqbal, Z., Khan, M.N., Lateef, M., 2000. Anthelmintic activity of medicinal plants with particular reference to their use in animals in the Indo-Pakistan subcontinent, Small Ruminant Research. https://doi.org/10.1016/S0921-4488(00)00163-2

Amal, M.N.A., Koh, C.B., Nurliyana, M., Suhaiba, M., Nor-Amalina, Z., Santha, S., Diyana-Nadhirah, K.P., Yusof, M.T., Ina-Salwany, M.Y., Zamri-Saad, M., 2018. A case of natural co-infection of Tilapia Lake Virus and Aeromonas veronii in a Malaysian red hybrid tilapia (Oreochromis niloticus × O. mossambicus) farm experiencing high mortality. Aquaculture 485, 12–16. https://doi.org/10.1016/j.aquaculture.2017.11.019

Bwalya, P., Simukoko, C., Hang’ombe, B.M., Støre, S.C., Støre, P., Gamil, A.A.A., Evensen, Ø., Mutoloki, S., 2020. Characterization of streptococcus-like bacteria from diseased Oreochromis niloticus farmed on Lake Kariba in Zambia. Aquaculture 523. https://doi.org/10.1016/j.aquaculture.2020.735185

Careaga, M. ónica, Fernández, E., Dorantes, L., Mota, L., Jaramillo, M.E., Hernandez-Sanchez, H., 2003. Antibacterial activity of Capsicum extract against Salmonella typhimurium and Pseudomonas aeruginosa inoculated in raw beef meat. Int. J. Food Microbiol. 83, 331–335. https://doi.org/10.1016/S0168-1605(02)00382-3

Cichewicz, R.H., Thorpe, P.A., 1996. The antimicrobial properties of chile peppers (Capsicum species) and their uses in Mayan medicine. J. Ethnopharmacol. 52, 61–70. https://doi.org/10.1016/0378-8741(96)01384-0

Confessor, M.V.A., Mendonça, L.E.T., Mourão, J.S., Alves, R.R.N., 2009. Animals to heal animals: Ethnoveterinary practices in semiarid region, Northeastern Brazil. J. Ethnobiol. Ethnomedicine 5. https://doi.org/10.1186/1746-4269-5-37

Cruz, P.M., Ibáñez, A.L., Monroy Hermosillo, O.A., Saad, H.C.R., 2012. Use of Probiotics in Aquaculture. Int. Sch. Res. Netw. ISRN Microbiol. 2012, 13. https://doi.org/10.5402/2012/916845

El-Deek, A.A., Al-Harthi, M.A., Osman, M., Al-Jassas, F., Nassar, R., 2012. Hot pepper (Capsicum Annum) as an alternative to oxytetracycline in broiler diets and effects on productive traits, meat quality, immunological responses and plasma lipids. Arch. Geflugelkunde 76, 73–80.

Fieira, C., Oliveira, F., Calegari, R.P., Machado, A., Coelho, A.R., 2013. Inibidores naturais no controle in vitro e in vivo de Penicillium expansum. Food Sci. Technol. 33, 40–46. https://doi.org/10.1590/S0101-20612013000500007

Guèye, E.F., 2002. Newcastle disease in family poultry: Prospects for its control through ethnoveterinary medicine, in: Livestock Research for Rural Development. pp. 80–91.

Hai, N.V., 2015. Research findings from the use of probiotics in tilapia aquaculture: A review, Fish and Shellfish Immunology. Academic Press. https://doi.org/10.1016/j.fsi.2015.05.026

Hanif, S.M., Meher, M.M., Biswas, G.C., Anower, A.M., 2012. FIELD STUDY ON EFFICACY OF RED PEPPER (Capsicum Annum) ALONG WITH ANTIBIOTICS AGAINST NEWCASTLE DISEASE IN BROILER AT NARAIL SADAR UPAZILLA, BANGLADESH. Wayamba J. Anim. Sci. 578, 1460–1466.

Harikrishnan, R., Balasundaram, C., 2005. Modern trends in Aeromonas hydrophila disease management with fish, Reviews in Fisheries Science. Taylor & Francis Group. https://doi.org/10.1080/10641260500320845

Khan, M.I.C.M.A., Hanif, W., 2006. Ethno veterinary medicinal uses of plants from Samahni valley Dist. Bhimber, (Azad Kashmir) Pakistan. Asian J. Plant Sci. 5, 390–396. https://doi.org/10.3923/ajps.2006.390.396

Kobata, K., Todo, T., Yazawa, S., Iwai, K., Watanabe, T., 1998. Novel Capsaicinoid-like Substances, Capsiate and Dihydrocapsiate, from the Fruits of a Nonpungent Cultivar, CH-19 Sweet, of Pepper (Capsicum annuum L.). J. Agric. Food Chem. 46, 1695–1697. https://doi.org/10.1021/jf980135c

Koffi-Nevry, R., Kouassi, K.C., Nanga, Z.Y., Koussémon, M., Loukou, G.Y., 2012. Antibacterial activity of two bell pepper extracts: Capsicum annuum L. and Capsicum frutescens. Int. J. Food Prop. 15, 961–971. https://doi.org/10.1080/10942912.2010.509896

Kurian, A.L., Starks, A.N., 2002. HPLC analysis of capsaicinoids extracted from whole orange habañero chili peppers. J. Food Sci. 67, 956–962. https://doi.org/10.1111/j.1365-2621.2002.tb09435.x

Marini, E., Magi, G., Mingoia, M., Pugnaloni, A., Facinelli, B., 2015. Antimicrobial and anti-virulence activity of capsaicin against erythromycin-resistant, cell-invasive group A streptococci. Front. Microbiol. 6. https://doi.org/10.3389/fmicb.2015.01281

Maroyi, A., 2012. Use of traditional veterinary medicine in Nhema communal area of the Midlands Province, Zimbabwe. Afr. J. Tradit. Complement. Altern. Med. 9, 315–322. https://doi.org/10.4314/ajtcam.v9i3.3

Omolo, A.M., Wong, Z.-Z., Mergen, A.K., Hastings, J.C., Le, N.C., Reiland, H.A., Case, K.A., Baumler, D.J., 2014. Antimicrobial Properties of Chili Peppers. J. Infect. Dis. Ther. 02, 145. https://doi.org/10.4172/2332-0877.1000145

Shen, S., Qian, J., Ren, J., 2010. Ethnoveterinary plant remedies used by Nu people in NW Yunnan of China. J. Ethnobiol. Ethnomedicine 6. https://doi.org/10.1186/1746-4269-6-24

Tran, N., Chu, L., Chan, C.Y., Genschick, S., Phillips, M.J., Kefi, A.S., 2019. Fish supply and demand for food security in Sub-Saharan Africa: An analysis of the Zambian fish sector. Mar. Policy. https://doi.org/10.1016/j.marpol.2018.11.009

Usman, M.G., Rafii, M.Y., Ismail, M.R., Malek, M.A., Latif, M.A., 2014. Capsaicin and dihydrocapsaicin determination in chili pepper genotypes using ultra-fast liquid chromatography. Molecules 19, 6474–6488. https://doi.org/10.3390/molecules19056474

Watts, J., Schreier, H., Lanska, L., Hale, M., 2017. The Rising Tide of Antimicrobial Resistance in Aquaculture: Sources, Sinks and Solutions. Mar. Drugs 15, 158. https://doi.org/10.3390/md15060158

